# Homology-based perspective on pangenome graphs

**DOI:** 10.64898/2026.03.16.712038

**Authors:** Anna Lisiecka, Agnieszka Kowalewska, Norbert Dojer

**Affiliations:** University of Warsaw; Ingenix.AI

## Abstract

Pangenome graphs conveniently represent genetic variation within a population. Several types of such graphs have been proposed, with varying properties and potential applications. Among them, variation graphs (VGs) seem best suited to replace reference genomes in sequencing data processing, while whole genome alignments (WGAs) are particularly practical for comparative genomics applications. For both models, no widely accepted optimization criteria for a graph representing a given set of genomes have been proposed.

In the current paper we introduce the concept of homology relation induced by a pangenome graph on the characters of represented genomic sequences and define such relations for both VG and WGA model. Then, we use this concept to propose homology-based metrics for comparing different graphs representing the same genome collection, and to formulate the desired properties of transformations between VG and WGA models. Moreover, we propose several such transformations and examine their properties on pangenome graph data. Finally, we provide implementations of these transformations in a package WGAtools, available at https://github.com/anialisiecka/WGAtools.

## 1 Introduction

*Pangenome graphs* combine reference genomes and their variants into coherent data structures, making them an intuitive way to represent genetic variation, from single nucleotide polymorphisms to large-scale structural variants (e.g. insertions, deletions and translocations). Technically, pangenome graphs have nodes labeled with sequences and are enhanced with collections of *genomic paths* representing individual genomes. Several types of such graphs have been proposed, differing in their properties, applications offered by available tools, efficiency, and even the level of precision with which the optimal graph representing given genomes is defined [3, 8]. Among them, variation graphs (VGs) and whole genome alignments (WGAs) are particularly widespread.

Each node of a VG is labeled with a fragment of one or more genomic sequences. The concatenated labels of successive nodes of a genomic path form the whole sequence of the represented genome. Such architecture enables efficient indexing methods [6, 16, 18, 22, 21], which is crucial for sequencing read mapping [11, 20]. Therefore, the VG model is probably the best pangenome model to replace reference genomes in processing DNA sequencing data.

WGA graphs (called Enredo graphs in [14]) have nodes labeled with multiple sequence alignments of homologous genome fragments [14, 19]. There is a one-to-one correspondence between aligned fragments and node occurrences in genomic paths; the latter define the order and orientation (DNA strand) of the fragments in the original sequences. The WGA graph structure naturally represents the structural variants found in individual genomes, while alignments complement this by showing the relationships between homologous genome fragments at the nucleotide level. Therefore, the WGA model is particularly useful for comparative genomics applications.

The basic idea underlying all pangenome graph models is that shared nodes along genomic paths indicate homologous genome regions. In de Bruijn graphs the decision when the paths share nodes is strictly determined by the length of the *k*-mers that form the nodes of the graph. In contrast, both VG and WGA models are highly flexible in the sense that the modeling framework leaves considerable freedom in deciding how homology and shared sequence segments are encoded in the structure. The strict criteria the structure of the variation graph should meet were proposed in [1] and implemented in Alfapang [2]. However, other widely used VG and WGA building tools heuristically infer the graph structure from pairwise genome alignments.

This raises the need to develop methods and tools for comparing and evaluating different graphs representing the same set of genomes. In the case of WGA, standards in this area were established by the Alignathon competition [7]. In the case of VG, simple graph characteristics (e.g., size) are typically compared, ignoring the way genomic sequences are represented within the graph. An exception is the approach proposed in [5], in which the metric is derived from the segmentations that genomic paths induce on genome sequences. Because segmentations are associated with homologies indicated in the graphs, this metric provides a better approximation of the similarity between the homologies.

In this article, we introduce the concept of a *homology relation*, which formally describes how a pangenome graph represents homology between genomic sequences. We then use this concept to propose a novel metric for comparing different VGs or WGAs representing the same set of genomes, specifically for evaluating a given graph against a gold standard. The proposed framework offers a unified perspective on both models (and potentially other pangenome models as well), paving the way for further applications: direct comparison of a VG and a WGA representing the same sequence set, as well as the transformations of one model to the other. We discuss the criteria that such transformations can/should meet and implement the proposed solutions in a package WGAtools. Finally, we evaluate the transformations and pangenome graph construction methods.

## 2 Preliminaries

### 2.1 Colinear alignment models

Consider a set of homologous sequences *S* = {*s*_1_, …, *s*_*n*_}. Multiple sequence alignment (MSA) of *S* is a matrix filled with sequence symbols in such a way that the concatenation of all non-empty entries from *i*-th row forms sequence *s*_*i*_.

Partial order alignment (POA) is a graph-based alternative to the matrix-based alignment representation. The set of nodes of the POA graph can be constructed from sequence residues by merging the ones that are located in the same MSA column and labeled with the same symbol (see Figure 1). The graph has two kinds of edges:

**Figure 1.**
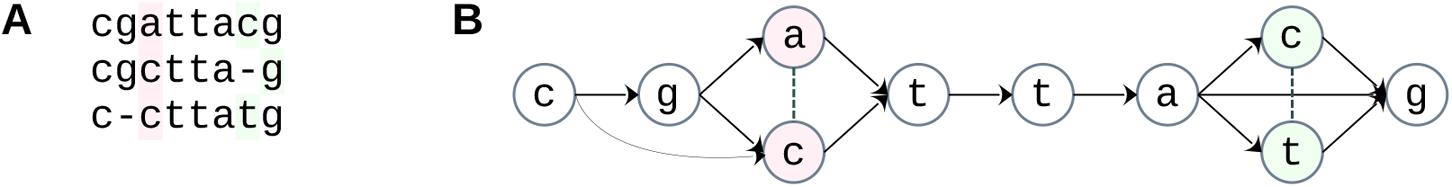
Matrix representation of MSA (**A**) and the corresponding POA graph (**B**).

- undirected edges that connect nodes representing aligned positions labeled with different symbols,
- directed edges that link nodes representing consecutive residues in *S*-sequences. The definition implies that the directed part of the POA graph is acyclic.

### 2.2 Non-colinear models and bidirected graphs

Neither MSA nor POA can represent homology between sequences affected by rearrangement events in their evolutionary history. In order to handle rearranged sequences, the models need to be generalized. For example, relaxing the acyclicity constraint would allow POA to represent sequences affected by translocations and duplications, but not inversions. The latter requires using pangenome models based on bidirected graphs.

In bidirected graphs, every node has two sides (denoted here ±1), and undirected edges connect adjacent nodes on specific sides. A node *v* can be *oriented* in two possible ways, denoted *v*^+1^and *v*^−1^. A *path* in a bidirected graph is a sequence of oriented nodes 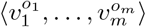such that for all respective *j*, side *o*_*j*−1_ of node *v*_*j*−1_ is connected by an edge to side −*o*_*j*_ of node *v*_*j*_ (so a path enters and exits the node on opposite sides). We refer to side −*o*_1_ of node *v*_1_ and side *o*_*m*_ of node *v*_*m*_ as *outer sides* of this path, while all other sides of its nodes are *inner* in this path. When all nodes in the path are different and each its inner side is incident with only one edge (connecting consecutive path nodes), the path is a *unitig*. A unitig having only one node is called *trivial*.

### 2.3 Variation graphs

Variation graph (VG) is a bidirected graph with a function *l* that labels nodes with DNA sequences. Two possible node orientations correspond to the two strands of DNA molecules. The labeling function extends to oriented nodes: *l*(*v*^+1^) = *l*(*v*) and *l*(*v*^−1^) is the reverse complement of *l*(*v*), and further to paths: 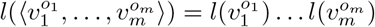.

A sequence *s* is *represented* in a VG by path *p* if *s* = *l*(*p*). In this case, *p* decomposes *s* into the labels of *p*-nodes or their reverse complements. Consequently, each nucleotide in *s* is represented by a specific nucleotide in the label of one of *p*-nodes. Moreover, both nucleotides are identical when the node has orientation +1 in *p* and are complementary otherwise.

When all sequences in *S* are represented in a VG by paths that jointly cover the whole graph, we say that the VG *represents S*, and the paths representing *S*-sequences are called *genomic paths*.

### 2.4 Whole genome alignments

Whole genome alignment (WGA) is a generalization of MSA. Instead of a single MSA of full input sequences, WGA consists of multiple MSAs that are built for sequence fragments or their reverse complements. The MSAs constituting a WGA are called *blocks*, and the sequences constituting a block are called *walks*. Walk has orientation −1 if it is a reverse complement of the input sequence fragment, and +1 otherwise. Usually it is assumed that the walks are disjoint, i.e. every sequence position is covered by at most one walk. Every WGA can be uniquely transformed into the one that satisfies the above condition by performing its *transitive closure* (implemented in `mafTransitiveClosure`tool of the `mafTools package`[7]). The number of walks in a block is called block *degree*. Blocks of degree 1 define no alignments between sequence fragments, so they can be added or removed from WGA without violating the represented genome alignment. In particular, every WGA can be supplemented with uncovered fragments of the input sequences transformed into single-walk blocks. Therefore, in the rest of the paper we assume without loss of generality that the WGA walks cover all positions of the sequences, and thus each sequence is a concatenation of its walks, appropriately ordered and oriented. This assumption allows us to define WGA graph as a bidirected graph, in which each sequence is represented as a path. Namely, the nodes of the graph are blocks, and side *o*_1_ of block *B*_1_ is connected by an edge to side *o*_2_ of block *B*_2_, if in some input sequence there are adjacent walks *w*_1_, *w*_2_ such that *w*_1_ occurs in *B*_1_ with orientation *o*_1_, and *w*_2_ occurs in *B*_2_ with orientation −*o*_2_.

## 3 Pangenome graph representation of sequence homology

### 3.1 Homology relations

Consider a set of homologous sequences *S* = {*s*_1_, …, *s*_*n*_} . We define the set of *positions* in *S* as *Pos*(*S*) = {⟨*i, j*⟩ |1 ≤*i* ≤*n*∧ 1 ≤*j* ≤|*s*_*i*_|}, where |*s*_*i*_| denotes the length of sequence *s*_*i*_. Every MSA (or POA) of *S* defines the following equivalence relation on *Pos*(*S*): positions ⟨ *i, j*⟩ and ⟨*i*′, *j*′⟩are *aligned* if and only if *s*_*i*′_[*j*′] andare placed in the same column of the MSA matrix (respectively are represented in the POA graph by the same node or by two nodes connected by an undirected edge).

The alignment relation indicates corresponding positions in *S*-sequences, describing their homology on the single-nucleotide level. Below we introduce similar *homology relations* associated with VGs and WGAs representing *S*. The definitions take into account that these models can represent homology between both parallel and antiparallel DNA strands.

Assume that positions ⟨*i, j*⟩, ⟨*i*′, *j*′⟩∈*Pos*(*S*) are covered by the same VG node shared by paths *p*_*i*_ and 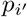 and represented by the same position in the label of this node. Then we say that ⟨*i, j*⟩ and⟨ *i*′, *j*′⟩ are:

- *merged directly* if this node has the same orientation on both paths,
- *merged inversely* if it has opposite orientations on each path.

Every WGA block defines the alignment relation on the positions of its walks. The following definition lifts all such relations to *Pos*(*S*), taking into account the orientation of the walks. Assume that positions

⟨*i, j*⟩ and ⟨*i*′, *j*′⟩ are covered by walks *w* and *w*′, respectively, of the same WGA block and are aligned in this block. Then we say that ⟨*i, j*⟩ and ⟨*i*′, *j*′⟩ are:

- *aligned directly* if *w* and *w*′have the same orientation,
- *aligned inversely* if *w* and *w*′have opposite orientations.

When positions ⟨*i, j*⟩ and ⟨*i*′, *j*′⟩are merged (aligned) directly or inversely, we say that they are simply *merged* (aligned, respectively). Note that the merging and direct merging relations are equivalence relations, and each equivalence class of the first one contains elements of one or two classes of the second one. The same applies to alignment and direct alignment relations, respectively.

### 3.2 Equivalent and canonical representations

We will say the two VG (WGA) representations of a sequence set *S* are *equivalent* if they induce the same homology relations on *Pos*(*S*), i.e. the same position pairs are merged (aligned, respectively) directly and the same pairs are merged (aligned, respectively) inversely by both graphs. To characterize the classes of equivalent representations, below we define modifications to the VG/WGA graph structure that do not change the induced homology relations.

*Splitting* a VG node *v* consists of replacing it with a unitig having |*l*(*v*) | nodes labeled with subsequent *l*(*v*) characters. Splitting operation can be reversed by *compressing* a unitig into a single node, labeled with the concatenation of the labels of the unitig nodes. Unitig is called *safe*, if no of its inner sides is an outer side of a genomic path (in particular, unitigs created with the splitting operation are safe).It is easily seen that operations of splitting a node and compressing a safe unitig transform a VG representation of a sequence set *S* into an equivalent representation of *S*.

Representation is *singular* if all its nodes are labeled with single characters (so it cannot be further split). Representation is *compact* if all its safe unitigs are trivial (so it cannot be further compressed). In Appendix A we prove that every class of equivalent VG representations has a singular and a compact representation and both have uniquely determined structures.

Similarly to VG nodes, WGA blocks may be split into single-column blocks without violating the alignment relations. However, in order to save the representations of sequence sets, subgraphs resulting from splitting blocks are not necessarily paths. The reason is that each gap in a walk requires an edge that skips columns filled with this gap (such edges are also added during the transformation of MSA into POA, see example in Figure 1).

Due to the above observation, in WGA the reverse operation has to be defined on acyclic subgraphs that are not necessarily paths. Although every such subgraph can be compressed into a single block, the structure of this block is not uniquely specified (see Figure 2 for example). Moreover, the maximal acyclic subgraphs may overlap, and, consequently, one WGA graph may be transformed to many non-isomorphic maximally compressed graphs.

**Figure 2.**
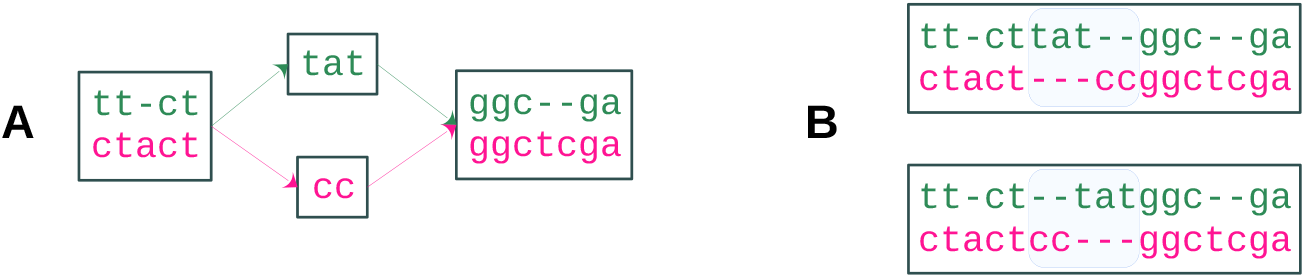
An acyclic WGA graph (A) that can be compressed into two different blocks (B).

### 3.3 Comparing representations

For both VG and WGA models there can exist multiple graphs representing the same collection of genome sequences. Representations could be compared based on graph characteristics (number of nodes, length of node labels, number of genomic paths covering the node, etc.), but, in general, the similarity of such characteristics poorly reflects the similarity of the represented sequence homologies. A better approximation is given by the *edit distance* [5], derived from the sequence segmentations induced by the genomic paths. However, this metric still does not allow direct comparison of the represented homologies, which can be done through the homology relations.

Two merging or two alignment relations may be compared using any set similarity measure, e.g. the Jaccard distance. Moreover, one can calculate the proportion of pairs of homologous positions in one of the compared relations among all pairs of homologous positions in the other relation. When the relation associated with the graph being analyzed is compared to the true one, these fractions define the *precision* and *recall* statistics, assessing how much the graph is biased by identifying nonexistent relationships or by missing existing ones.

The above metrics allow comparison of graph representations up to equivalence. In the case of VG, equivalent representations differ only in the granularity of the graph. The compact representation provides maximum compression, while the singular one ensures that each fragment of the input sequences is represented by a complete path. In the case of WGA, the criteria for selecting the best representation in a class of equivalent representations are not as clear. Although WGA with single-column blocks can serve as a canonical representation, such blocks are not what one can expect from good alignments. Indeed, representations with blocks containing large genomic fragments with high homology are more informative. Therefore, more compressed graphs are preferred, and blocks should be split into shorter ones only when the homology level for one or more walks decreases.

The homology level within blocks can be measured using the two complementary metrics: the proportion of matching nucleotides within the alignment relation and the fraction of gaps in walks. The former (called *identity score* in [17]) is determined by the alignment relation and thus is constant throughout a class of equivalent representations. On the other hand, the fraction of gaps can be used as an indicator of whether the blocks are excessively compressed.

Some of the above statistics can be calculated for WGAs using mafTools package [7]. We have implemented the remaining ones as part of the WGAtools package, available at https://github.com/anialisiecka/WGAtools. All the statistics can be also calculated for VGs, after converting it to WGA using the `vg2wga` tool (described in the next section), which is included in WGAtools as well.

## 4 Model transformations

Given set of sequences *S*, both VG and WGA representation of *S* define homology relations on the same set *Pos*(*S*), but these relations are subject to different constraints. Two positions can be merged directly (inversely) only when the nucleotides on these positions are identical (respectively complementary). On the other hand, any two positions can be aligned either directly or inversely. Therefore, WGA can fully represent sequence homology at the nucleotide level, whereas VG only represents homology of matching nucleotides and ignores mismatched ones. Hence, we will say that a WGA representation of *S* is *compatible* with a VG representation of *S*, when any two positions ⟨*i, j*, ⟩ ⟨*i*′, *j*′⟩∈*Pos*(*S*) are merged directly (inversely) in VG if and only if they are aligned directly (respectively inversely) in WGA and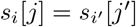 (respectively *s*_*i*_[*j*] and 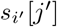 are complementary).

Transformations between WGA and VG can be expected to ensure compatibility between the input and output representations. In the case of transformation from WGA to VG, this condition actually determines the structure of the resulting VGs up to equivalence, the corresponding algorithm is presented in section 4.1. On the other hand, there does not seem to be a canonical way to perform the inverse transformation. Design decisions may depend on the answers to the following questions:

- should the transformation focus on compatibility or on inferring homology between mismatched nucleotides, missed in VG?
- how to select a proper balance between graph compactness (large blocks) and low gap content within the blocks (which is easier to obtain with short blocks)?
- how to choose VG subgraphs to be transformed into single blocks when the candidate subgraphs overlap?

In section 4.2, we present three algorithms that implement solutions based on different answers to the above questions. The novel transformation algorithms are included in the WGAtools package, available at https://github.com/anialisiecka/WGAtools.

### 4.1 Transformation of WGAs into VGs

Our transformation algorithm, called `wga2vg,` consists of three steps:

1. **Transforming WGA blocks into POA graphs**. Each block is transformed into a POA graph, which is further considered as a special case of a VG graph – undirected edges are ignored, directed ones connect sides +1 of their origins with sides −1 of their targets.
2. **Transforming WGA edges into VG edges**. Edges joining blocks in a WGA graph are transformed into VG edges joining nodes representing end positions in respective walks of these blocks. Thus the graph gains edges connecting nodes representing consecutive positions in input sequences that belonged to different WGA blocks.
3. **Compacting the graph**. Maximal safe unitigs are compressed into single nodes. This operation reduces the size of the graph without violating the merging relations on input sequences.

Figure 3 visualizes the algorithm on a toy example. In Appendix A we prove that wga2vg produces a compact VG representation that is compatible with the input WGA representation.

**Figure 3.**
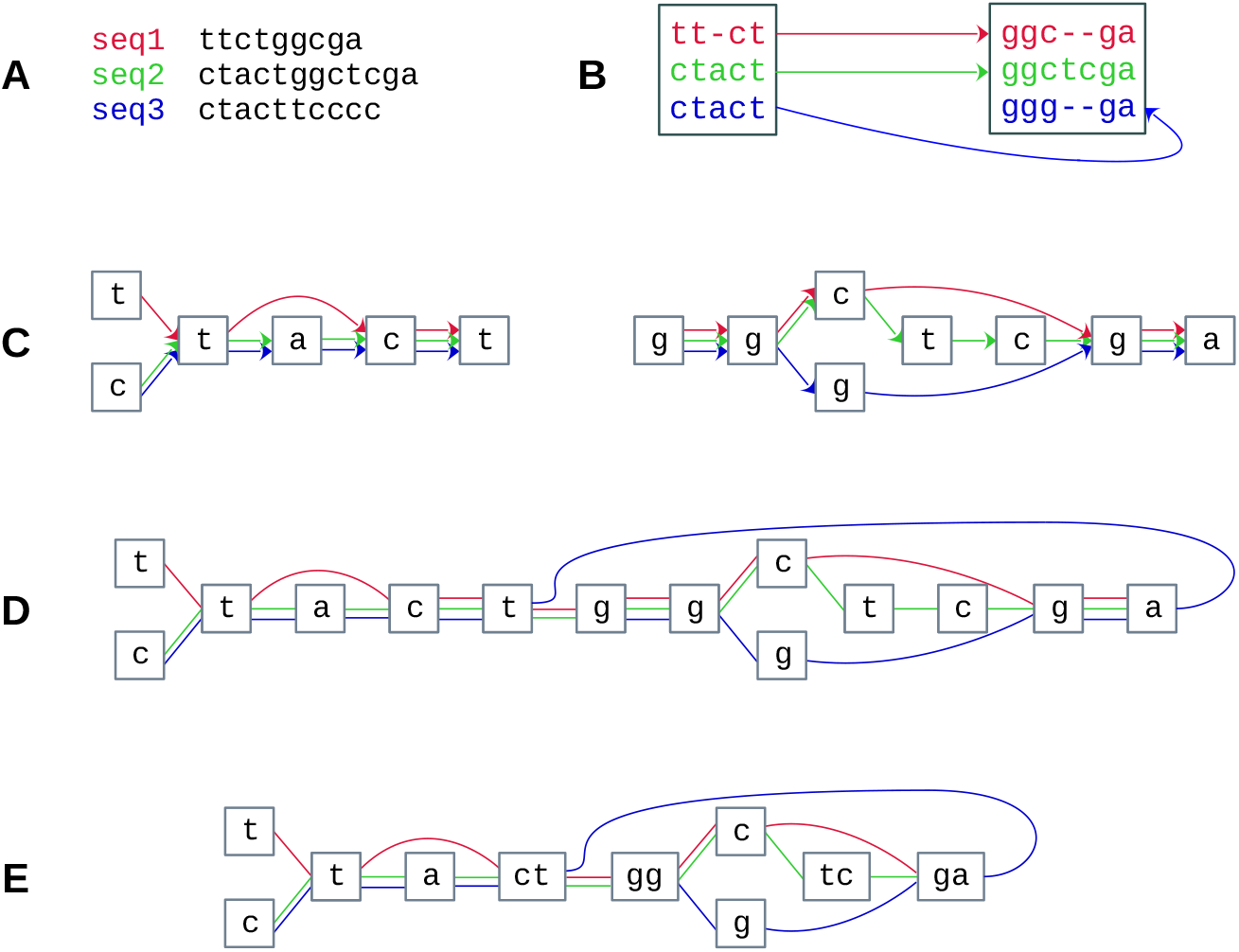
Transformation of a whole genome alignment graph into a variation graph. **A-B** Input DNA sequences and their whole genome alignment representation. Each edge represents the adjacency of the incident block labels in one of the input sequences. The blue sequence in the right block is the reverse complement of the second half of sequence seq3. **C** POA graph representations of the WGA blocks. **D** Variation graph after including edges between the WGA blocks. **E** The final graph with unbranched paths compacted into nodes.

### 4.2 Transformations of VGs into WGAs

Below we describe three different algorithms transforming VGs into WGAs, implemented in the maffer tool by Garrison and Seumos [10] and in two novel tools: `vg2wga` and `block-detector`. The approaches are illustrated in Figure 4.

**Figure 4.**
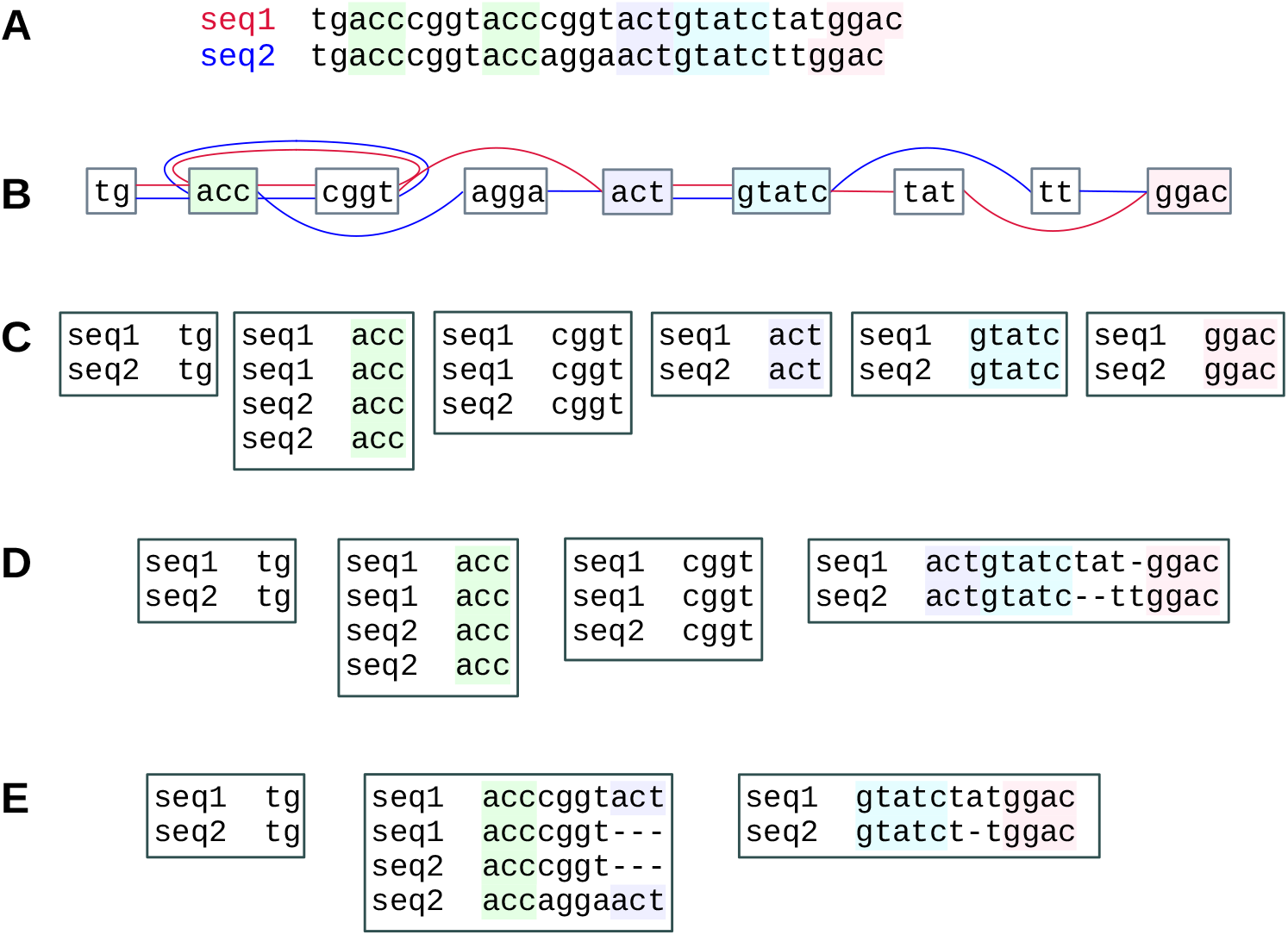
Different transformations of a variation graph into a whole genome alignment graph. **A-B** Input DNA sequences and their variation graph representation. **C-E** Resulting WGAs (blocks containing only one segment are omitted). **C** Whole genome alignment generated by `vg2wga`. Blocks correspond to the nodes of the variation graph. **D** Whole genome alignment generated by `maffer`. The first three blocks are exactly the same as in `vg2wga,`the last one corresponds to the fragment of VG between the purple and the pink node. **E** Whole genome alignment generated by `block-detector`. The first block is the same as in `vg2wga` and `maffer,`the other two are build from VG nodes between the green and the purple node, and between the blue and the pink node, respectively.

In WGA graphs constructed using `vg2wga`, blocks correspond to the nodes of the input VGs. More precisely, each node is transformed into a block consisting of walks corresponding to the occurrences of this node in the genomic paths (see Figure 4C). Thus, `vg2wga` ensures compatibility between the representations and yields WGAs with no gaps in blocks and with identity score equal to 1. Consequently, the statistics listed in section 3.3 can be calculated for variation graphs by applying WGA analysis tools (such as mafTools) to the WGAs outputted by the `vg2wga` transformation. Since `vg2wga` does not infer homology between mismatched nucleotides (i.e. leaves them unaligned), resulting WGAs may be not suitable for some applications.

The second tool, `maffer,` assumes that the nodes of an input variation graph have been *linearized* (i.e. sorted and oriented) in a reasonable way. The ordered list of input VG nodes is split into intervals inducing topologically sorted subgraphs that are further transformed into separate WGA blocks (see Figure 4D). Thus, `maffer` can create significantly more complex blocks than`wga2v g` does, but the result depends on the way the nodes have been linearized. Optimization criteria and algorithms solving the linearization problem were discussed in several works (see e.g. [15]). Authors of `maffer` recommend to accomplish this task using the `odgi sort` tool [12], so in the rest of this paper we assume these programs are used together as a pipeline.

The third algorithm, `block-detector,` searches the variation graph to identify subgraphs that are suitable for transformation into a single WGA block (see Figure 4E). The searching algorithm is inspired by the one used in whole-genome aligner SibeliaZ [17]. SibeliaZ determines the walks forming the resulting WGA block using a de Bruijn graph built from genomic sequences. The main idea is that the paths representing walks in this graph form a subgraph containing a so called *carrying path* that has densely distributed common fragments with each of the paths representing walks. `block-detector` identifies analogous subgraphs in variation graphs using an algorithm very similar to the one of SibeliaZ. Then, the walks are aligned using common path fragments as alignment anchors. The details of the `block-detector` algorithm are described in the Appendix B.

## 5 Results

We conducted three series of experiments. First, we compared VGs constructed using different tools from the same input sequences. Then, we compared the properties of the VG-to-WGA transformations. Finally, we evaluated the ability of the VG-building tools combined with VG-to-WGA transformations to reconstruct the alignment relation corresponding to the simulated evolutionary history.

In all these experiments we used alignments previously used for evaluation of SibeliaZ [17]. Alignments were generated with ALF [4], a tool to simulate bacterial genome evolution. The evolutionary model of ALF includes both point mutations (insertions, deletions and substitutions) and large-scale structural variants (e.g., insertions, translocations and gene gain/loss). ALF generates an alignment corresponding to the evolutionary history of genomes, in our experiments used as the *ground truth*. We used six datasets, simulated with different phylogenetic distances from genomes to their common ancestor, ranging from 0.03 to 0.18 substitutions per site. Each dataset consists of 10 bacterial genomes of size around 1.5Mbp, composed of 1500 genes.

Each dataset served as input to the following VG-building tools: PanGenome Graph Builder (PGGB) [9], Minigraph-Cactus [13], AlfaPang and AlfaPang+ [2]. PGGB constructs VG in three steps: all-to-all genome alignments, initial graph inference from the alignments and graph refinement. Minigraph-Cactus constructs an initial graph representing large structural variants by iterative genome alignment and then completes it to a full variation graph. The structure of the final graph depends on a predetermined reference genome, which is guaranteed to be represented by an acyclic path. AlfaPang uses alignment-free and reference-free approach, relying on intrinsic sequence properties. Finally, AlfaPang+ is the pipeline consisting of AlfaPang and the graph refinement step of PGGB.

In the first experiment, all these tools were applied to genomic sequences generated by ALF and the output VGs were compared using two metrics: the Jaccard distance and the normalized version of the edit distance proposed in [5]. The original edit distance was defined in [5] as the number of sequence breakpoints induced by only one of the two graphs. We divide this number by the total number of breakpoints induced by both graphs, which ensures that the distance falls in the range [0,1], as the Jaccard distance does. Figure 5 summarizes the results. For each pair of VG-building tools, both metrics follow the same trend – the distance increases with increasing divergence between input sequences. Moreover, both metrics have very similar values for the comparison of graphs constructed using PGGB, Minigraph-Cactus and AlfaPang+. On the other hand, when comparing a graph created using AlfaPang with a graph from another tool, the edit distance is much larger than the Jaccard distance, especially for datasets with low genomic divergence. The reason is that repetitive sequence regions are represented in graphs created using AlfaPang by complex local structures, which induce numerous breakpoints. Other tools avoid such structures by forcing the acyclicity of the path representing the reference genome (Minigraph-Cactus) or by “smoothing” the graph in the refinement step (PGGB and AlfaPang+). The impact of the refinement on both metrics is particularly well illustrated in the comparison of distances between graphs created with AlfaPang and AlfaPang+ .

**Figure 5.**
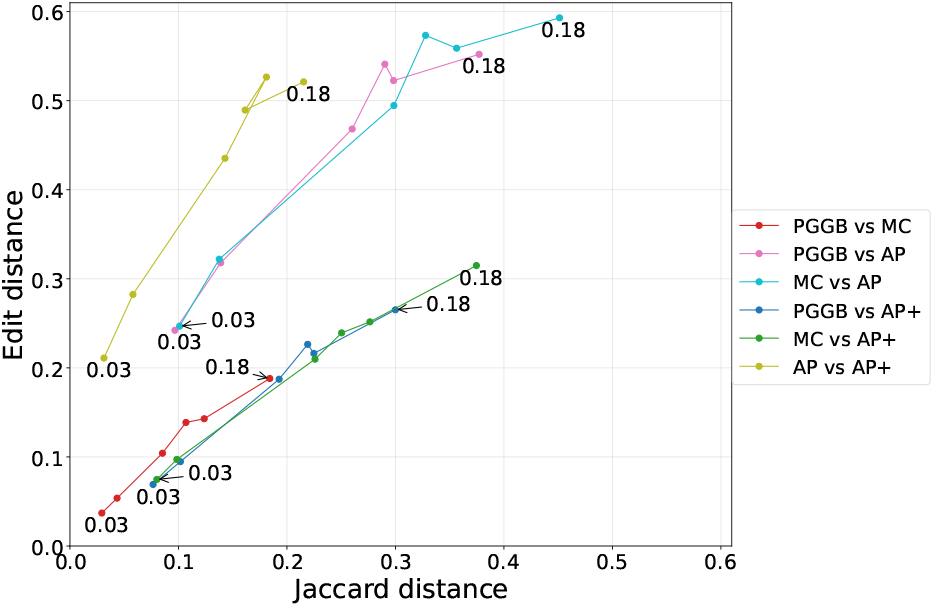
Distances between different variation graphs. Numerical labels correspond to the genomic divergence of the simulated dataset. MC: Minigraph-Cactus, AP: AlfaPang, AP+: AlfaPang+

In the second experiment, we evaluated the ability of the VG-to-WGA transformations to recover the alignment relation from the merging relation. To this aim, WGAs outputted by ALF were first transformed to VGs using `wga2vg` and then transformed back to WGAs using `vg2wga`, `maffer` and `block-detector`. Figure 6 summarizes the performance of these tools and the properties of the resulting WGAs. As expected, `vg2wga` is the most efficient. Both `vg2wga` and `maffer` are significantly faster than `block-detector` and consume more than twice less memory. `block-detector` produces the least fragmented alignments. It has 1-2 orders of magnitude less blocks and longer sequences than `maffer`, which in turn has an order of magnitude less blocks and longer sequences than`vg2wga`. Hence, `vg2wga` produces WGA graphs with very large number of blocks (100k-450k), consisting of very short sequences (4-15bp on average). The lowest average block degree have alignments produced using `vg2wga` and the highest – the ones produced using `maffer`. `vg2wga` creates alignments with full identity score and no gaps. The lowest average identity and the largest fraction of gaps have WGAs outputted by `maffer`, which may explain how this tool achieves the highest genome coverage by blocks. In almost all cases, `block-detector` algorithm has the closest statistics to the ground truth.

**Figure 6.**
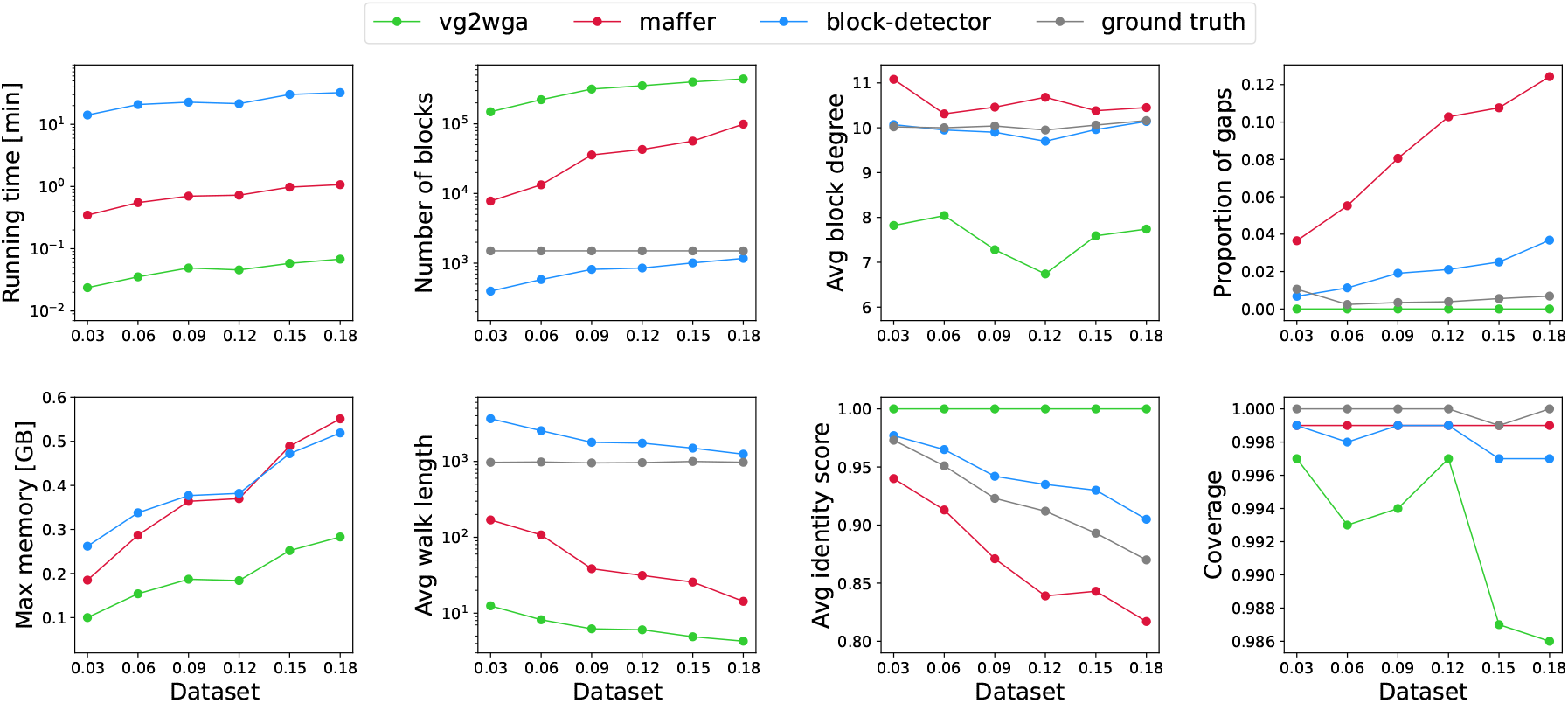
Efficiency and properties of WGA graphs obtained from VGs using different transformation methods. In each plot, numerical labels on x-axis correspond to the genomic divergence of the dataset.

In the third experiment, we applied the VG-to-WGA transformations to VGs constructed from genomic sequences using different tools. In terms of the characteristics shown in Figure 6, the resulting WGAs do not differ significantly from those obtained in the previous series of experiments (the detailed comparison is presented in Appendix C). Figure 7 shows the accuracy of the reconstruction of the ground truth WGA. The results obtained for the VGs transformed from WGAs using `wga2vg` may serve as a reference point. Due to its design principles, `vg2wga` gets always precision 100%, and recall depending on the phylogenetic distance between the genomes (here in the range 94–99%). `maffer` and `block-detector` have slightly lower precision (above 99.7%) and higher recall (above 97% for `maffe</monosr` and 99.6% for `block-detector`). In the case of VGs built directly from genomic sequences, the precision still has very high values (above 0.98% across all datasets), but much wider spread of recall results is observed. For PGGB and Minigraph-Cactus it varies between 58% and 92%. This is visibly lower than for AlfaPang (above 75%) and much lower than for AlfaPang+ (above 90%). For all the VG building tools and datasets `vg2wga` has the highest precision among all transformation tools, while `block-detector` – the highest recall. Moreover, in most cases `block-detector` has higher precision than `maffer`.

**Figure 7.**
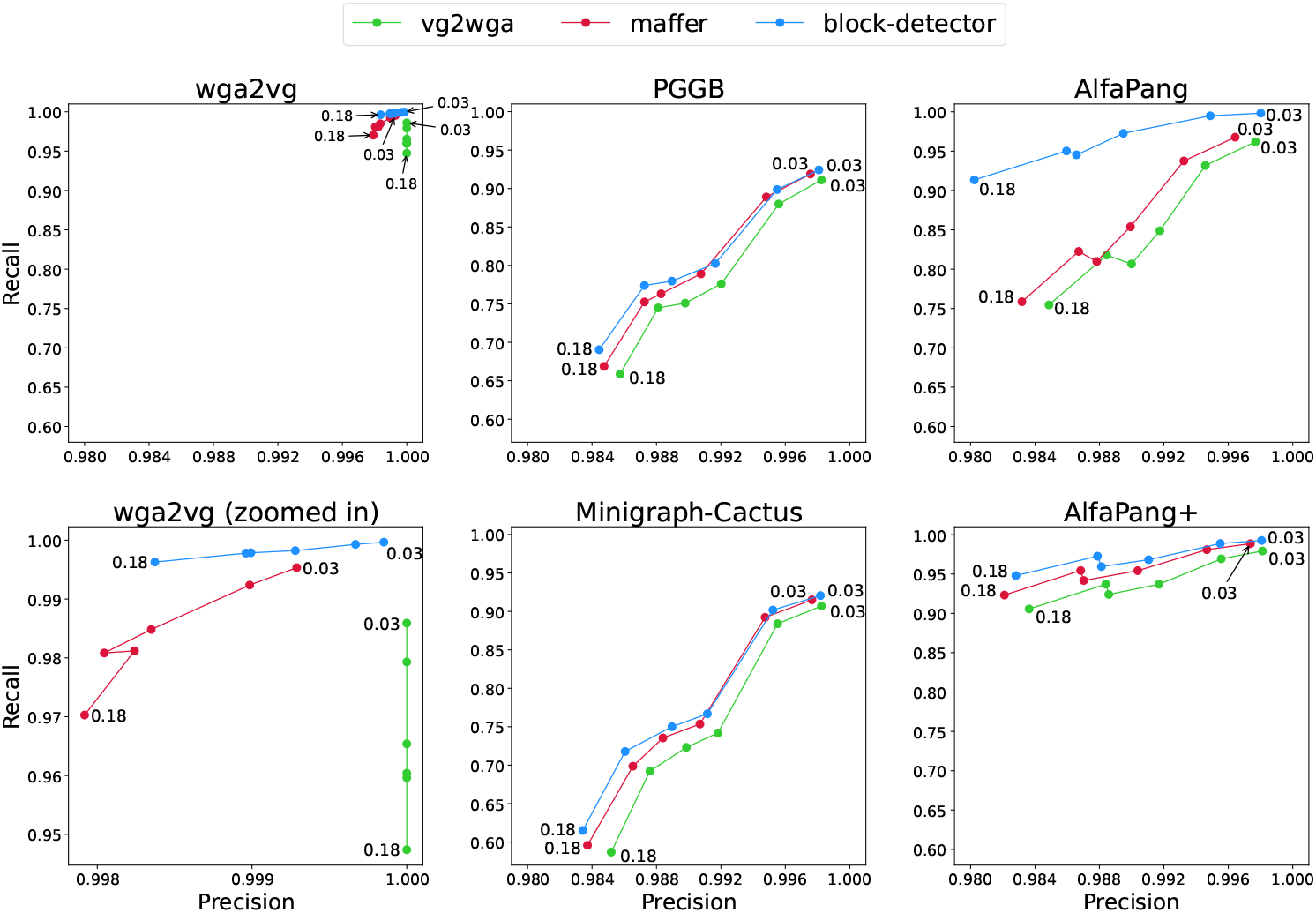
Alignment reconstruction accuracy for VGs transformed from ground true WGA (left column) and for VGs built from genomic sequences only (middle and right column). In each plot, the values corresponding to the least and most divergent datasets are labeled by their genomic divergence.

**Figure 8.**
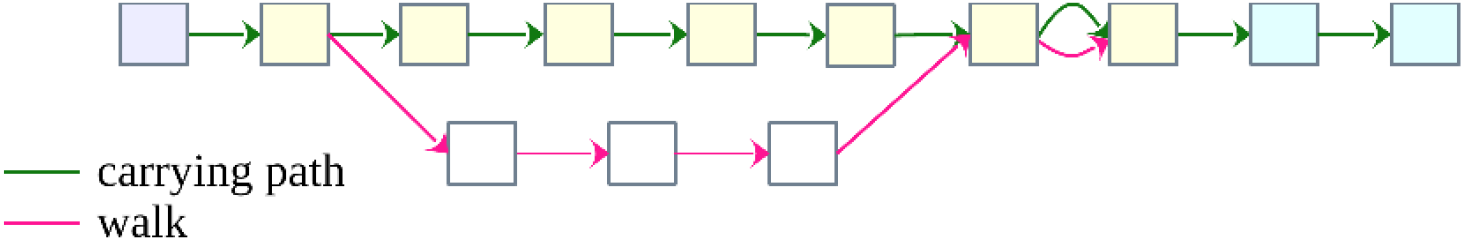
Variation graph representing a walk (pink edges) and a carrying path *q* (green edges) of a block. The purple, yellow and blue nodes belong to *q*_1_, *q*_2_, *q*_3_ of the partition of *q*, respectively.

## 6 Conclusion

We have proposed a framework for comparing pangenome graph representations of a genome collection. The framework is based on the concept of representation-induced homology relation, which can be defined for different pangenome models. Our contributions include:

- homology-based metrics for comparing representations both within and across models,
- analysis of the structure of classes of VG and WGA representations sharing the same relation,
- design and implementation of transformations between VG and WGA representations: `wga2vg, vg2wga and block-detector,`
- evaluation of transformations from VG to WGA representations.

In the context of homology relations, the problem of transforming a WGA representation into a VG representation has a straightforward canonical solution, implemented in our tool `wga2vg:` the input relation is restricted to matching nucleotides. On the other hand, inverse transformations can extend the homology relation to mismatched nucleotides in a variety of ways. Our evaluation shows that:

- `vg2wga,` which does not infer homology between mismatched nucleotides, is extremely fast and memory-efficient, but generates highly fragmented WGAs,
- `block-detector` is computationally demanding, but yields the highest inference accuracy (precision and recall *>* 99% in our experiments),
- Garrison and Seumos’s `maffer` represents a compromise between computational efficiency and inference accuracy, although its output contains excessive gaps.

We also investigated WGA construction pipelines, consisting of VG building tools and VG-to-WGA transformations. Our experiments show that the first step of the pipeline has greater impact on the accuracy (especially the recall) of the resulting WGAs than the transformation step. The best results (recall >95% and precision >98%) were obtained for the pipeline consisting of AlfaPang+ and `block-detector`.

## Acknowledgements

This work was supported by the National Science Centre, Poland, under grant number 2022/47/B/ST6/03154.

## A Theorems and proofs

### A.1 Canonical VG representations

Two bidirected graphs are *isomorphic* if there exist

- a bijection *i* between the nodes of the graphs,
- an orientation change indicator function *c* assigning each node ±1,

such that nodes *v*_1_ and *v*_2_ of the first graph are connected by an edge on sides *o*_1_ and *o*_2_, respectively, iff *i*(*v*_1_) and *i*(*v*_2_) are connected by an edge on sides *c*(*v*_1_) · *o*_1_ and *c*(*v*_2_) · *o*_2_, respectively.

#### Theorem 1.

*Equivalent singular VG representations of a sequence collection are isomorphic*.

*Proof*. In a singular representation each node corresponds to exactly one equivalence class of its merging relation. Since equivalent representations of a sequence set *S* induce the same merging relations on *Pos*(*S*), the above correspondence establishes the bijection *i* between the sets of nodes in any two equivalent singular VGs. Moreover, *i* satisfies the property that for every node *v* in the first graph, the labels *l*(*v*) and *l*(*i*(*v*)) are either identical or complementary. In the first case we set *c*(*v*) = +1, otherwise *c*(*v*) = −1.

We will show that the functions *i* and *c* establish the isomorphism between the graphs. To this aim choose nodes *v*_1_ and *v*_2_ of the first graph that are connected by an edge on sides *o*_1_ and *o*_2_, respectively. Since all edges must be covered by genomic paths, either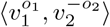 or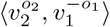 is a subpath of some genomic path, and thus *v*_1_ and *v*_2_ represent neighboring positions in the respective *S*-sequence. In the other graph these positions are represented by nodes *i*(*v*_1_) and *i*(*v*_2_), which must be connected by an edge on sides *c*(*v*_1_) · *o*_1_ and *c*(*v*_2_) · *o*_2_, respectively. Similar argument applies to the opposite implication.

#### Theorem 2.

*Equivalent compact VG representations of a sequence collection are isomorphic*.

*Proof*. We first show that two different maximal safe unitigs cannot share an oriented node. To this aim assume that two such unitigs have one. By definition, it cannot be followed nor preceded by two different nodes in these two unitigs. Since the unitigs are different, the end node of one unitig must be an inner node of the other one. Thus the path forming the first unitig can be extended at this end with the next node from the other unitig. This contradicts the maximality of the first unitig, unless the node extending the path already belongs to it. However, in this case the unitig forms a maximal non-cyclic path on a cycle, in which every node is incident only to the edges of that cycle. Finally, the cycle must be covered by genomic paths and thus must contain an outer side of a genomic path, which contradicts the assumption that both unitigs are safe.

The consequence of what we have shown is that two maximal safe unitigs that share a node are either identical or reversed paths. Therefore the maximal safe unitigs divide the set of VG nodes into subsets that can be compressed independently. Thus every compact representation is the result of compressing all maximal safe unitigs in an equivalent singular representation. When two compact representations are equivalent, the singular representations are also equivalent and hence isomorphic. Consequently, equivalent compact representations are isomorphic too.

### A.2 Properties of the wga2vg transformation

#### Theorem 3.

*The transformation* `wga2vg` *produces a compact VG representation that is compatible with the input representation*.

*Proof*. The merging relation is established in step 1 of the algorithm and further steps leave it unmodified. Compatibility follows directly from the definition of POA graph. Step 2 ensures that each input sequence is represented in the graph as a single path. Finally, step 3 makes the representation compact.

## B Details of the block-detector algorithm

### B.1 VG subgraphs representing blocks

The notation introduced in this section is based on [17], but the definitions are adapted to VGs.

A subpath of a genomic path is called *genomic walk*. A set of genomic walks *B* forms a *block* if the subgraph induced by *B* contains a *carrying path* that has densely distributed common fragments with each walk of *B*. The distance between consecutive common fragments is controlled by parameter *b* (by default *b* = 200). Below we provide a formal definition of a block and its score.

Two paths 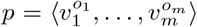 and 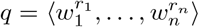form a *b*-*bubble*, if the following conditions are satisfied:

- 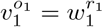,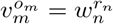and there are no other common nodes in paths *p* and *q*,
- |*l*(*p*)| ≤ *b* + |*l*(*v*_1_)| + |*l*(*v*_*m*_)| and |*l*(*q*)| ≤ *b* + |*l*(*w*_1_)| + |*l*(*w*_*n*_)|.

Two paths form a *b*-*chain*, if their common nodes form subsequences splitting them into *b*-bubbles. More formally, given paths 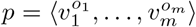 and 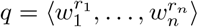, there should exist sequences of indexes 1 = *i*_1_ *<* … *< i*_*k*_ = *m* and 1 = *j*_1_ *<* … *< j*_*k*_ = *n* satisfying the following conditions:

- 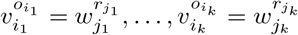 and there are no other common nodes in *p* and *q*,
- subpaths 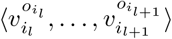 and 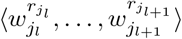 form a *b*-bubble for each *l*.

A set of genomic walks forms a *block B* with a *carrying path q*, if for each walk *p* ∈*B*, the carrying path can be split into subpaths *q* = *q*_1_*q*_2_*q*_3_ satisfying the conditions:

- *q*_2_ forms a *b*-chain with *p*,
- |*l*(*q*_1_)| ≤ *b* and |*l*(*q*_3_)| ≤ *b*.

The score of *p* with respect to *q* is then defined as *f* (*p, q*) = |*l*(*p*) | −(| *l*(*q*_1_)|+ |*l*(*q*_3_) |). Finally, the score of a block *B* with respect to carrying path *q* is given by formula

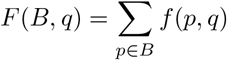

### B.2 Blocks finding algorithm

`block-detector` constructs blocks in a seed-and-extend manner. It starts with an arbitrary node (called *seed*) and tries to extend it into a carrying path that induces a block consisting of walks corresponding to homologous sequences. After the seed is chosen, the block is initialized with all the occurrences of the seed on genomic paths that do not belong to any block yet.

The elongation process is performed in several steps. Each step starts with computing the carrying path extension. To this aim, `block-detector` computes for each genomic walk *p* its *b*-*extension*, i.e. the longest genomic walk *p*_*e*_ = *pq* such that | *l*(*q*) |≤ *b*. Next, the node *t*, to which the carrying path will be extended, is selected such that the number of occurrences of *t* in all *b*-extensions is maximized. Then, the carrying path is elongated by the shortest fragment *r* of a *b*-extension, linking the current end of the carrying path with node *t*. The process is illustrated in Figure 9. After extending the carrying path by a walk *r*, the remaining walks of a block are elongated. The elongation of walks is held in steps, each taking into consideration a single node *v* of *r*. For each unused occurrence *occ* of *v*, the algorithm checks if there is a walk whose *b*-extension contains *occ*. If such a walk does not exist, a new walk is started using just *occ*. Otherwise, the found walk is elongated by its *b*-extension, truncated at *occ*. If there are multiple walks found, the one which ends further in the genome is chosen. The elongation step ends with updating the *b*-extensions of all walks. If the score of the extended block is non-negative, the elongation process is continued. Otherwise, the process is finished and the extension of the seed with the highest score is recalled and saved into the final result.

**Figure 9.**
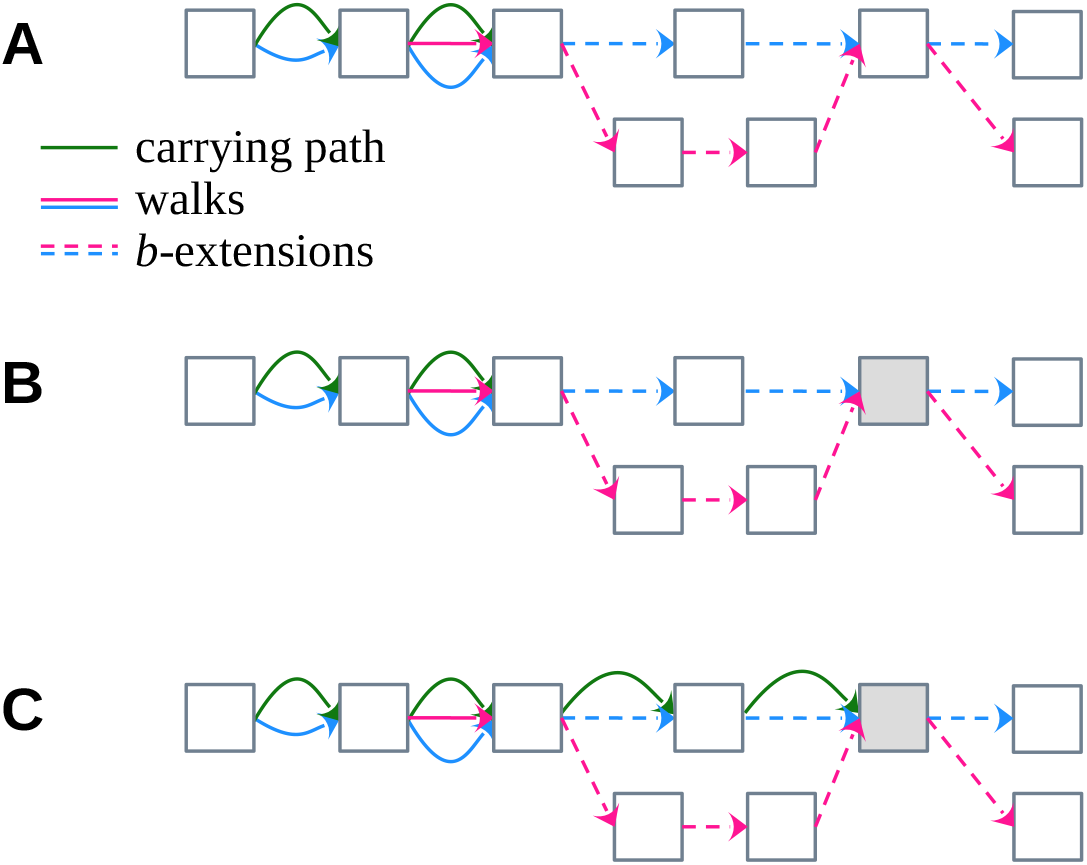
Carrying path elongation. **A** Graph with a block consisting of two walks (pink and blue) and an initial fragment of the carrying path (green). The *b*-extensions of the walks are marked with dashed lines. **B** Grey vertex is the one chosen as the future end of the carrying path. **C** Carrying path extended till the grey vertex.

The main pseudo-code of `block-detector` is shown in Algorithm 1.

#### Algorithm 1 Blocks-finding algorithm

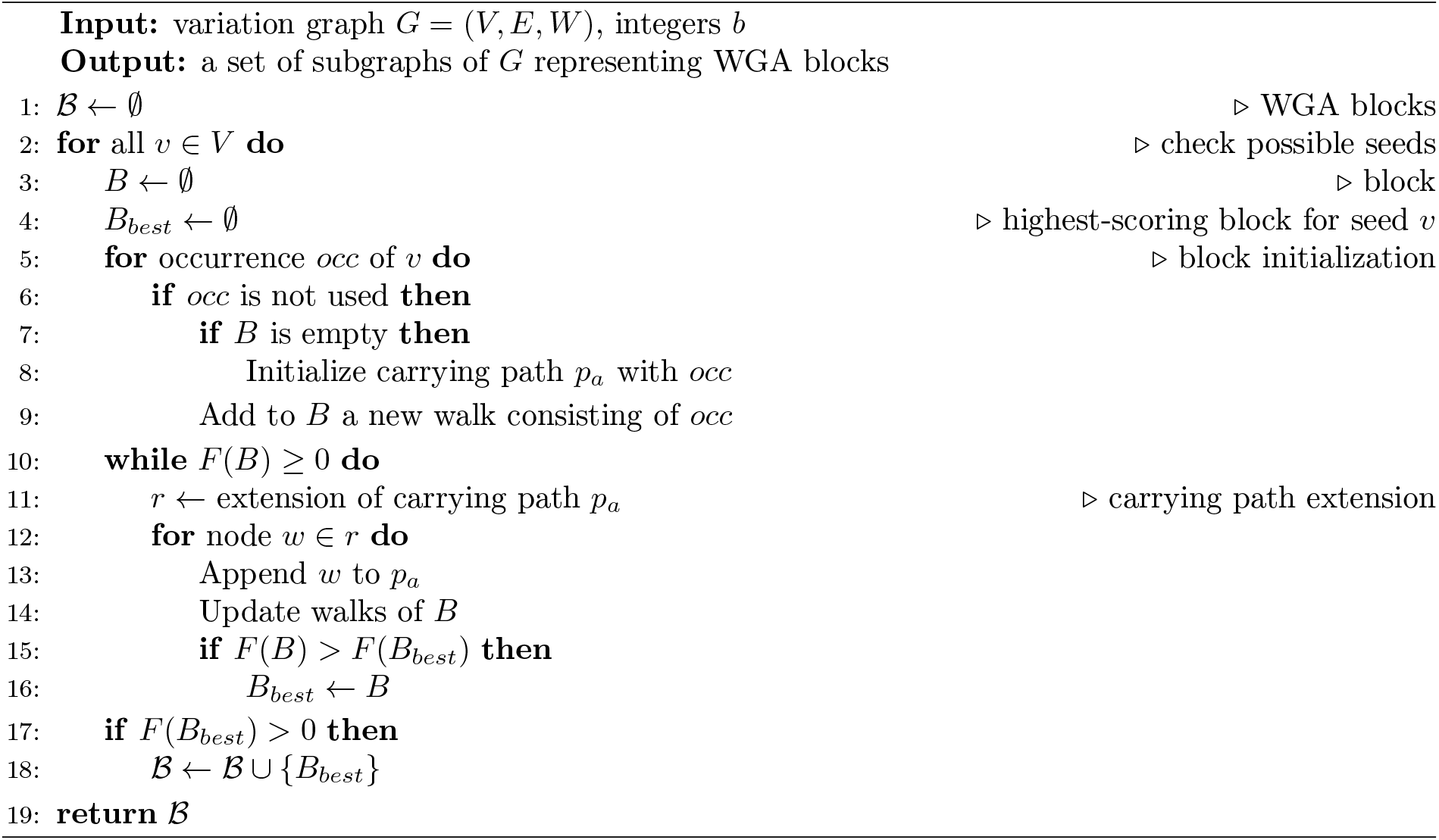

### B.3 Multiple sequence alignment

The alignment step in `block-detector` is based on the classic POA algorithm, modified to use the structure of the input VG subgraph as the skeleton of the final alignment. The POA graph is initialized with the carrying path of a block. Then, walks are aligned to the POA graph one at a time. Fragments of a walk represented in VG by a common path with the carrying path are aligned directly to the corresponding vertices of the POA graph. These alignments act as anchors for aligning the rest of the path. Namely, the subpath connecting two consecutive anchors in the path is aligned to the subgraph located between the anchors in the POA graph. The modified alignment is illustrated in Figure 10, on a simple example of one walk.

**Figure 10.**
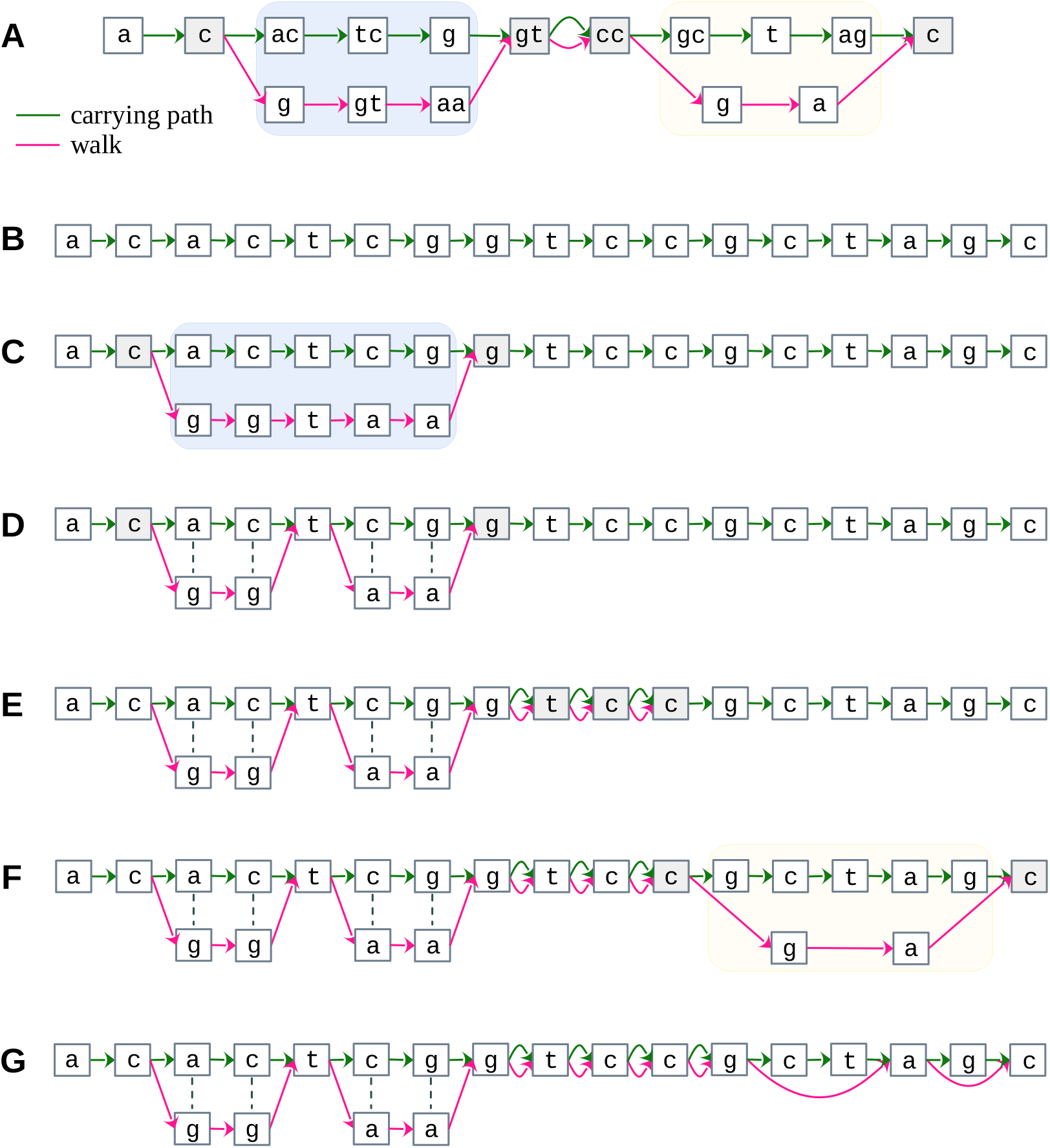
Consecutive steps of aligning a walk to a POA graph. **A** VG representing a walk (pink) and a carrying path (green). Their common nodes are colored in grey. The only two non-trivial bubbles, marked in blue and yellow (without their endpoints), represent the only fragments of the sequences that need to be aligned. **B** The first step of the alignment – POA graph initialized with the carrying path. **C** POA graph representing the carrying path along with the fragment (marked in blue) of the walk forming a bubble with the carrying path in VG. **D** POA graph after aligning the fragment of the sequence between the two common grey vertices. Mismatches are marked with dashed lines. **E** POA graph after mapping the aligned sequence on the three common grey vertices. **F** POA graph with added fragment of the walk, forming a bubble with the carrying path in VG (marked in yellow). **G** The final POA graph representing the alignment of the whole walk and the carrying path.

### B.4 Remarks

The algorithm of `block-detector` is adapted from SibeliaZ [17] with several modifications. Firstly, `block-detector` takes as input a variation graph instead of a compacted de Bruijn graph. In order to simplify the design of the graph structure, `block-detector` operates on nodes instead of edges, e.g. nodes are used as the seeds of blocks. Another difference concerns the block elongation process of both algorithms. In SibeliaZ, the *b*-extensions of walks are computed before the carrying path elongation and are unchanged during the walk elongation process. On the other hand, `block-detector` updates the *b*-extension of each walk after its elongation steps and uses it for computing the extension of the carrying path.

Probably the most important difference concerns the computation of the block alignments. In SibeliaZ walks are aligned from scratch using the SPOA tool [23]. In contrary, the alignment algorithm of `block-detector` uses the structure of the input VG subgraph as the skeleton of the final alignment. As a result, the common fragments of the carrying path and the paths representing walks are transformed into common paths of the POA graphs, thanks to which the transformation guarantees the preservation of homology between these fragments.

## C Supplementary results

Below we compare the characteristics of WGAs obtained from VGs created using different algorithms.

Figure 11 shows basic statistics regarding WGA blocks: number, average length and degree. In most cases, the charts for different VG building algorithms are quite similar to the ones for VGs obtained using the `wga2vg` transformation. The only exception is Minigraph-Cactus, for which all the transformations achieve relatively low average block degree, what can be partially explained by the way Minigraph-Cactus constructs VGs and by difficulties to find paralogous sequences in these graphs.

**Figure 11.**
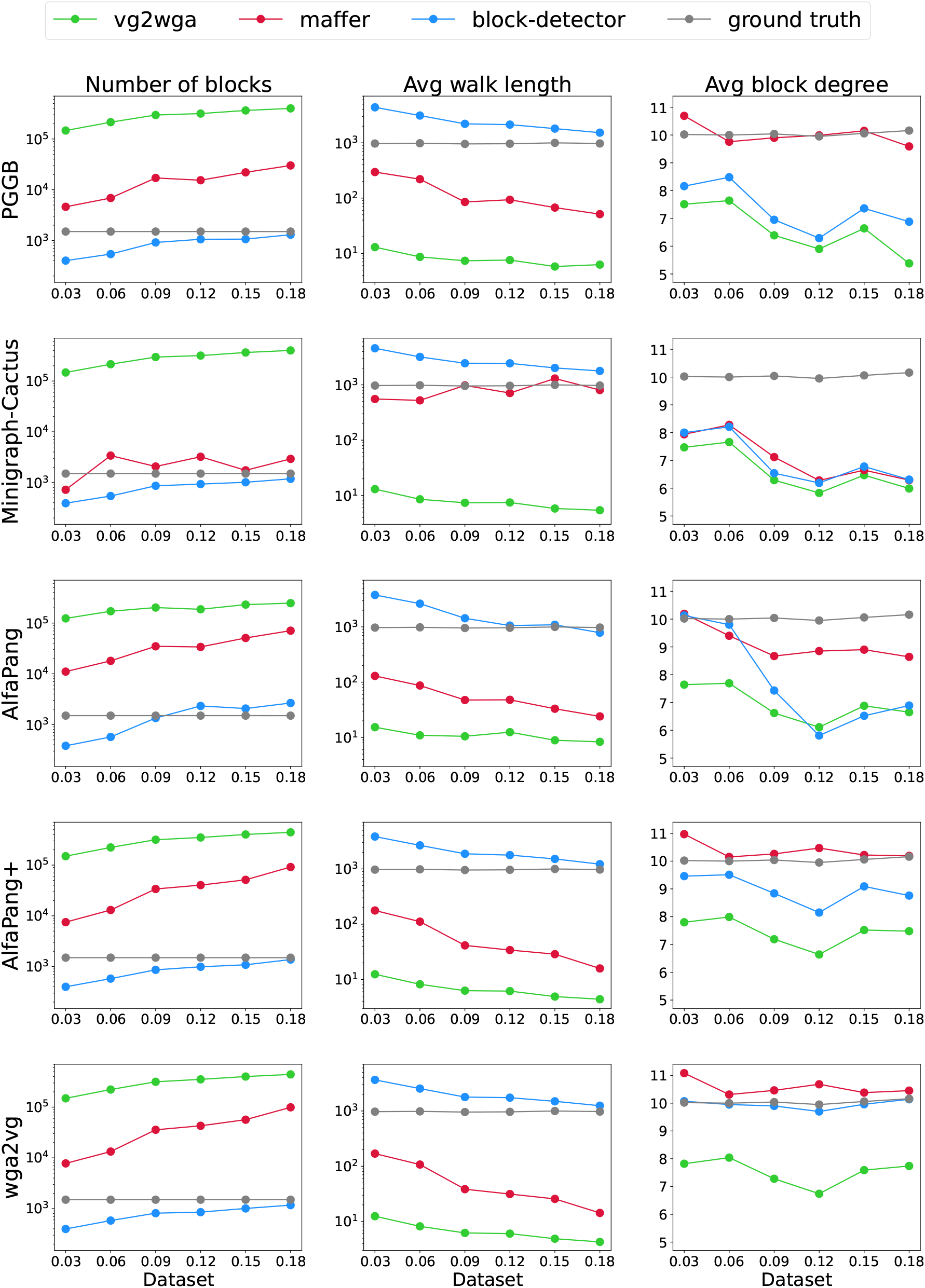
Number of blocks, their average length and degree in WGA graphs obtained using different VG building algorithm and transformation method. On the x-axis of each plot, bacterial datasets are labeled by their genomic divergence.

Figure 12 shows the coverage, average column identity score and fraction of gaps in the alignment blocks. For all graphs, `maffer` achieves the highest coverage values (above 0.99), while `vg2wga` – the lowest. block-detector achieves coverage above 0.97 for AlfaPang, AlfaPang+ and `wga2vg,` while for PGGB and Minigraph-Cactus its coverage is only slightly higher than those of `vg2wga,` and up to 0.1 lower than those of `maffer`. The average column identity score is consistently larger for block-detector than for `maffer` across all graphs. For PGGB, Minigraph-Cactus and AlfaPang graphs, the average column identity score of `maffer` is relatively low even for the least divergent bacterial dataset and drops by more than half for most divergent datasets. These values are partially explained by fraction of gaps in the alignments. For these graphs, the alignments generated by `maffer` consist of 10–60% of gaps compared to at most 7% for `block-detector`. For other graphs, going from least to most divergent bacterial dataset, the average column identity of `block-detector` drops from 0.98 to 0.90, while for `maffer` these value drops from 0.94 to 0.79.

**Figure 12.**
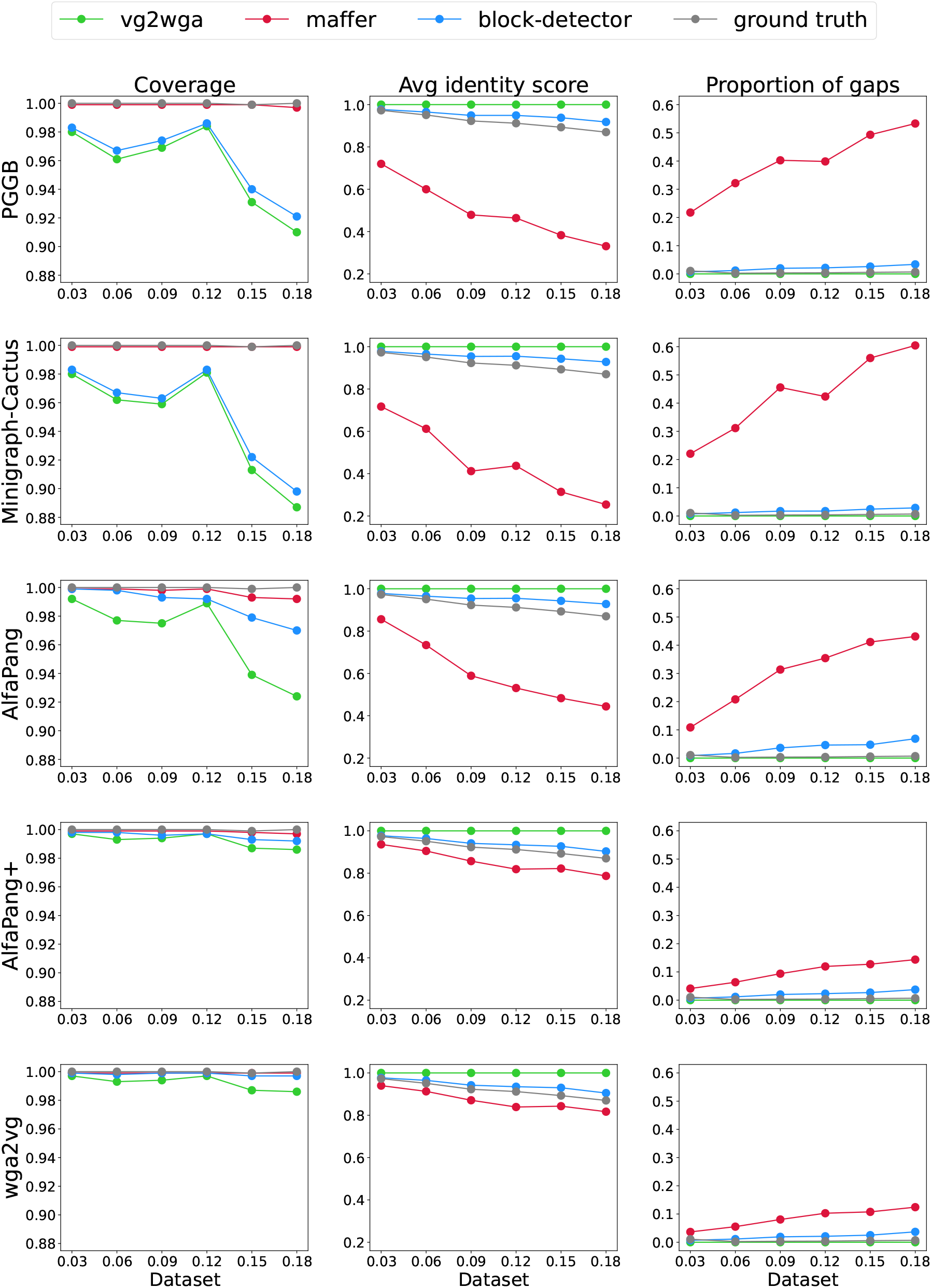
Coverage, average column identity and fraction of gaps in WGA graphs obtained using different VG building algorithm and transformation method. One-degree blocks were omitted in the calculations. On the x-axis of each plot, bacterial datasets are labeled by their genomic divergence.

## D Parameter and dataset details

Both PGGB and Minigraph-Cactus were run with default parameters. AlfaPang and AlfaPang+ were run with *k* = 25, and in the refinement step of the latter `smoothxg` and `gfaffix` were run with their default parameters. The simulated data were obtained from [17]. The link to the simulation parameter files, the simulated genomes, and their alignments is available at https://github.com/medvedevgroup/SibeliaZ/blob/master/DATA.txt.

